# Structural insights into the orthosteric inhibition of P2X receptors by non-ATP-analog antagonists

**DOI:** 10.1101/2023.09.15.558037

**Authors:** Danqi Sheng, Chenxi Yue, Fei Jin, Yao Wang, Muneyoshi Ichikawa, Ye Yu, Chang-Run Guo, Motoyuki Hattori

**Affiliations:** State Key Laboratory of Genetic Engineering, Shanghai Key Laboratory of Bioactive Small Molecules, Collaborative Innovation Center of Genetics and Development, Department of Physiology and Biophysics, School of Life Sciences, Fudan University, 2005 Songhu Road, Yangpu District, Shanghai 200438, China; School of Basic Medicine and Clinical Pharmacy, China Pharmaceutical University, Medical Building, Room 128, 639 Long-Mian Road, Nanjing 200098, China; State Key Laboratory of Genetic Engineering, Department of Biochemistry and Biophysics, School of Life Sciences, Fudan University, Shanghai 200438, China

## Abstract

P2X receptors are extracellular ATP-gated ion channels that form homo-or heterotrimers and consist of seven subtypes. They are expressed in various tissues, including neuronal and nonneuronal cells, and play critical roles in physiological processes such as neurotransmission, inflammation, pain, and cancer. As a result, P2X receptors have attracted considerable interest as drug targets, and various competitive inhibitors have been developed. However, although several P2X receptor structures from different subtypes have been reported, the limited structural information of P2X receptors in complex with competitive antagonists hampers the understanding of orthosteric inhibition, hindering the further design and optimization of those antagonists for drug discovery.

Here, we determined the cryo-EM structures of the mammalian P2X7 receptor in complex with two classical competitive antagonists of pyridoxal-5’-phosphate derivatives, PPNDS and PPADS, at 3.3 and 3.6 Å resolution, respectively, and performed structure-based mutational analysis by patch-clamp recording as well as MD simulations. Our structures revealed the orthosteric site for PPADS/PPNDS, and structural comparison with the previously reported apo-and ATP-bound structures showed how PPADS/PPNDS binding inhibits the conformational changes associated with channel activation. In addition, structure-based mutational analysis identified key residues involved in the PPNDS sensitivity of P2X1 and P2X3, which are known to have higher affinity for PPADS/PPNDS than other P2X subtypes. Overall, our work provides structural insights into the orthosteric inhibition and subtype specificity of P2X receptors by the classical P2X antagonists, pyridoxal-5’-phosphate derivatives, thereby facilitating the rational design of novel competitive antagonists for P2X receptors.

## Introduction

ATP not only serves as a cellular energy currency but also plays a key role in signal transmission for cellular stimulation between cell surface receptors^1^. P2X receptors are the family of cation channels activated by extracellular ATP and are widely expressed in the mammalian nervous, respiratory, reproductive, and immune systems^2^ ^3^ ^4^ ^5^. There are seven subtypes (P2X1-P2X7) in the family, each of which plays distinct roles in physiological and pathophysiological functions via homo-or heterotrimerization^6, 7^. In recent years, there has been growing interest in the development of drugs targeting the P2X family due to its involvement in various physiological and pathological conditions, and various antagonists have been developed^8^. Some have progressed to clinical trials^9^, and Gefapixant, a P2X3 receptor antagonist for chronic cough, was already on the market after the clinical study^10^.

ATP analogs are most common among competitive inhibitors for P2X receptors; however, they are generally unsuitable for in vivo applications due to their relatively low specificity, which may result in off-target toxicity. This issue arises because the human body contains numerous ATP-binding proteins. Therefore, a non-ATP-analog P2X inhibitor would be a more promising target to develop and optimize, and pyridoxal phosphate-6-azophenyl-2′,5′-disulfonic acid (PPADS) and its analog pyridoxal-5’-phosphate-6-(2’-naphthylazo-6’-nitro-4’,8’-disulfonate) (PPNDS) are such classical non-ATP-analog P2X inhibitors, namely, pyridoxal phosphate derivatives^11, 12, 13, 14^. PPNDS and PPADS belong to the class of competitive antagonists that selectively inhibit P2X receptors^15^, and P2X receptors are known to exhibit variable sensitivity to PPADS/PPNDS depending on the species and the specific subtype^14, 16, 17, 18, 19, 20^. It is noteworthy that P2X1 and P2X3 receptors show relatively high sensitivity to PPADS, but P2X2 and P2X7 receptors show only moderate sensitivity, and P2X4 receptors are insensitive to PPADS^21^.

Several attempts have been made to optimize pyridoxal phosphate derivatives as P2X antagonists^22, 23, 24, 25^. For example, the introduction of bulky aromatic groups at the carbon linker in PPADS was attempted to improve in the subtype specificity profiles, potentially opening up new avenues for targeted drug development and therapeutic intervention^23^. Despite the recent significant increase in structural information on P2X receptors^26, 27, 28, 29, 30, 31, 32, 33, 34^, the lack of structural information on P2X receptors in complex with pyridoxal phosphate derivative inhibitors has hampered their rational optimization for drug discovery targeting P2X receptors.

In this work, we determined the cryogenic electron microscopy (cryo-EM) structures of the panda P2X7 receptor in complex with PPADS and PPNDS. The structures revealed the orthosteric binding site for these pyridoxal phosphate derivatives. Structural comparison with the previously determined apo and ATP-bound P2X7 receptors^31, 34^ showed PPADS/PPNDS-dependent structural rearrangement at the orthosteric binding site for channel inactivation. Further mutational analysis by electrophysiological recording identified key residues of human P2X1 and P2X3 that show high sensitivity to pyridoxal phosphate derivative inhibitors.

## Results

### Structural determination and functional characterization

To gain insight into the mechanism of P2X receptor inhibition by pyridoxal phosphate derivatives, we used giant panda (*Ailuropoda melanoleuca*) P2X7, whose structures in complex with allosteric modulators have been reported^29^. Notably, panda P2X7 shares 85% identity with the human P2X7 receptor and exhibits high and stable expression profiles suitable for structural studies^29^. We performed whole-cell patch clamp recordings using HEK293 cells transfected with full-length panda P2X7 (pdP2X7). The application of 10 µM PPNDS and 100 µM PPADS blocked approximately 50% of the ATP-dependent currents from pdP2X7 **(Fig. S1, A-B and E-F**). This result correlates well with the properties of P2X7 receptors moderately inhibited by PPADS/PPNDS^17, 35^.

We then expressed and purified the previously reported crystallization construct of pdP2X7_cryst 29_. The purified pdP2X7_cryst_ was reconstituted into lipid nanodiscs, mixed with PPNDS and PPADS, separately, and subsequently subjected to single-particle cryo-EM (**Fig. S2-S5)**. The structures of pdP2X7 in the presence of PPNDS and PPADS were determined at 3.3 Å and 3.6 Å, respectively (**Table S1**).

The overall structures are similar and show the trimeric architecture of P2X receptors, consisting of the extracellular domain and two transmembrane (TM) helices, with each protomer resembling the dolphin shape, consistent with the previously reported P2X structures^36^ (**Fig. 1 and Fig. S6, A-B**). More importantly, we identified the residual EM densities at the agonist binding site that fit into the shape of PPNDS and PPADS (**Fig. 1**). It should be noted that while the pyridoxal phosphate groups in PPNDS and PPADS are shared, the naphthylazo group of PPNDS is significantly bulkier than the azophenyl group of PPADS. This difference facilitated our assignment of compound binding poses corresponding to each EM density (**Fig. 1**). Consistent with antagonist binding, the TM domain adapts to the closed conformation of the channel, as similarly observed in the previously reported closed state structures of P2X7 receptors^29, 34^.

**Figure. 1.**
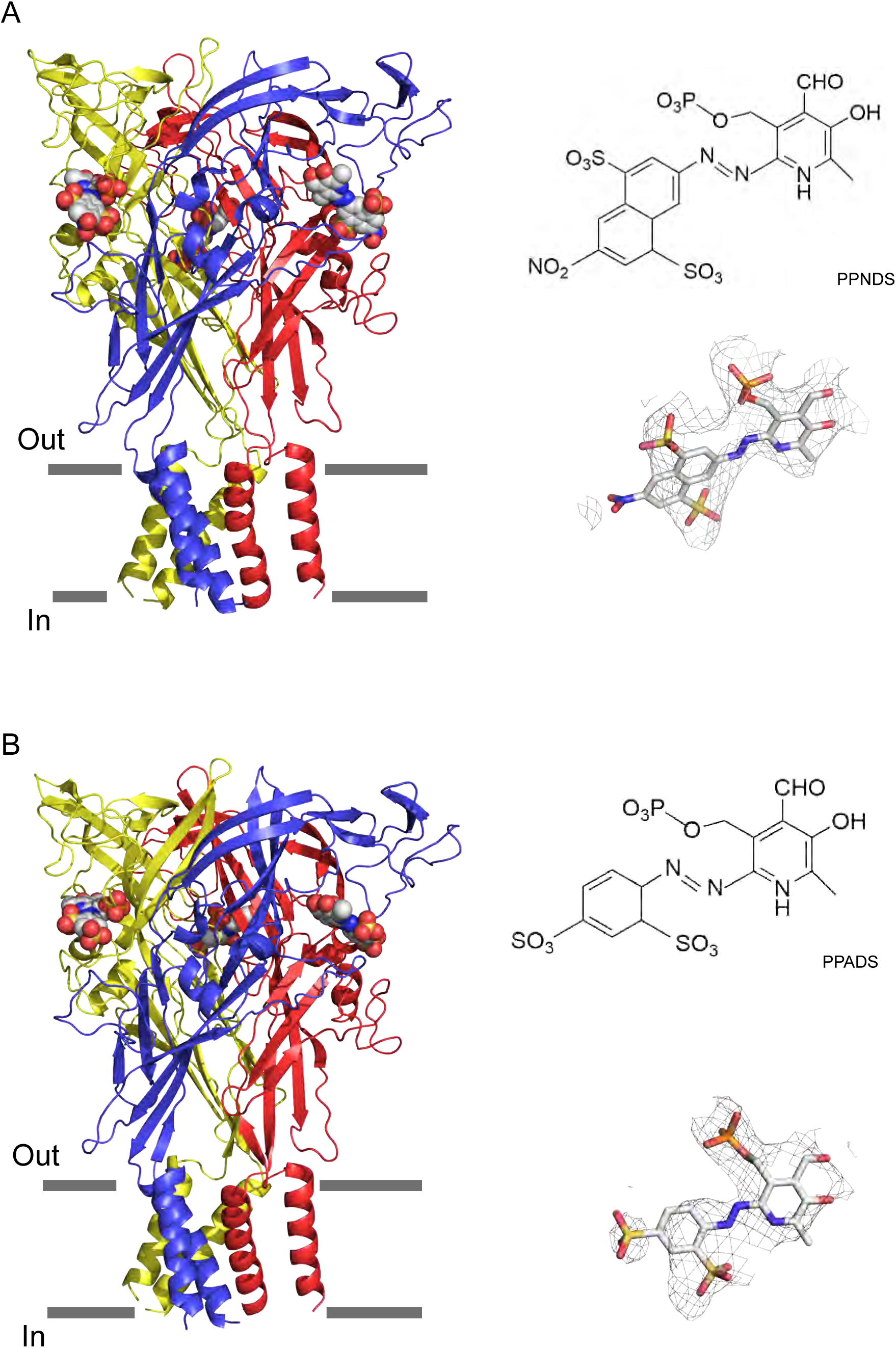
Cryo-EM structures of PPNDS-bound and PPADS-bound pdP2X7. The trimeric structures of PPNDS-bound (A) and PPADS-bound (B) pdP2X7, viewed parallel to the membrane. The PPNDS and PPADS molecules are shown as spheres. Each subunit of the trimers is colored blue, yellow, and red. The EM density maps contoured at 4.5 σ and 3.5 σ for PPNDS and PPADS are shown as gray mesh. The structural formulas of PPNDS and PPADS are also shown.

### Orthosteric binding site

In the PPNDS-bound and PPADS-bound structures, PPNDS and PPADS molecules bind to essentially the same orthosteric site, consistent with both PPNDS and PPADS being pyridoxal phosphate derivatives (**Fig. 2 and 3 and Fig. S7**). Furthermore, the residues involved in PPNDS and PPADS largely overlap with the residues at the ATP binding site in the previously reported ATP-bound P2X7 structure (**Fig. 4**)^34^. Many of the residues are highly conserved among P2X receptors and have been shown to be crucial for P2X activation^27, 37, 38, 39^, which is consistent with both of them being competitive inhibitors.

**Figure. 2.**
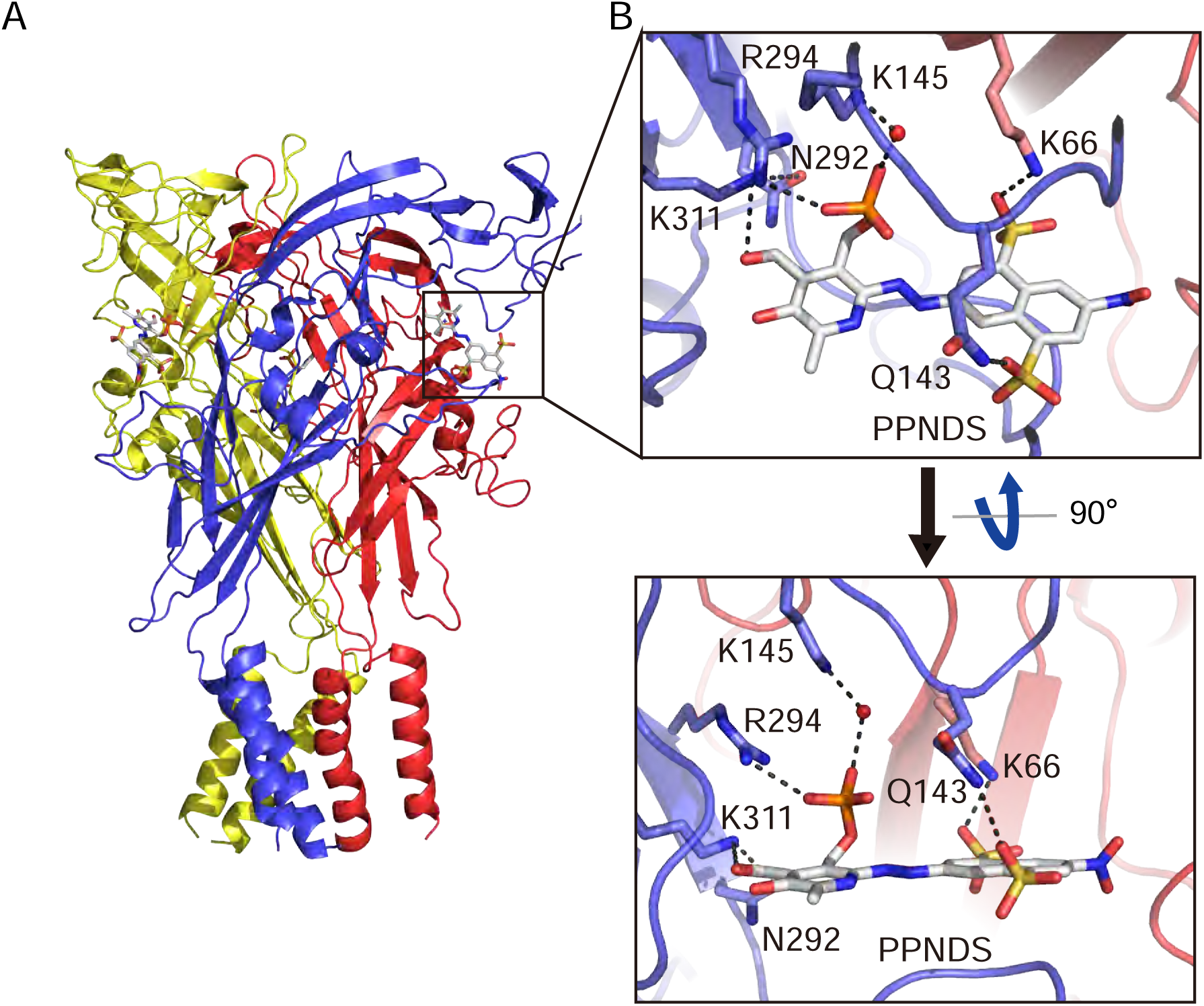
Binding site for PPNDS. (A, B) Overall structure (A) and close-up view of the PPNDS binding site (B) in the PPNDS-bound pdP2X7 structure. PPNDS molecules are shown by stick models. Water molecules are depicted as red spheres. Dotted black lines indicate hydrogen bonding.

**Figure. 3.**
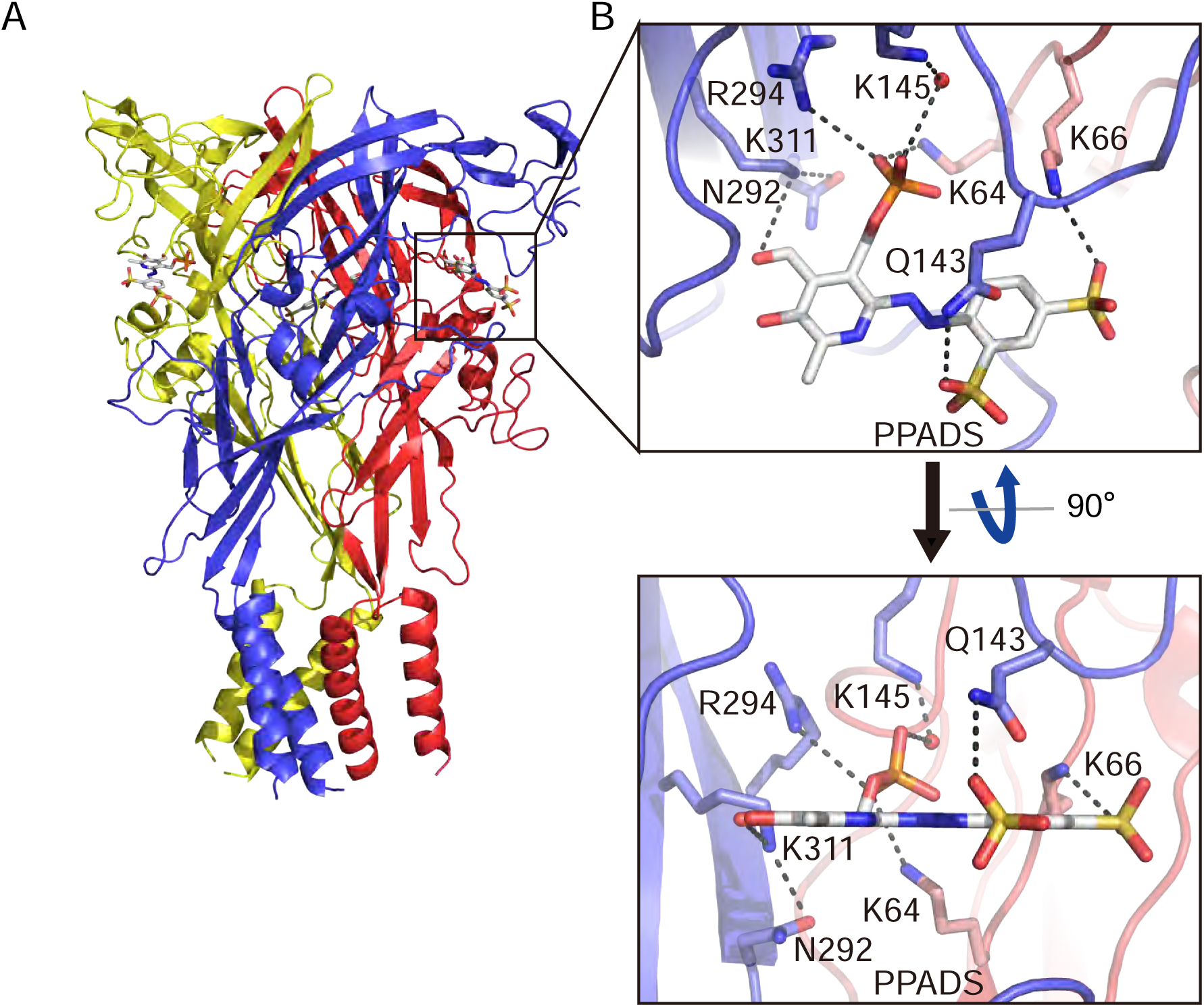
Binding site for PPADS. (A, B) Overall structure (C) and close-up view of the PPADS binding site (D) in the PPADS-bound pdP2X7 structure. PPADS molecules are shown by stick models. Water molecules are depicted as red spheres. Dotted black lines indicate hydrogen bonding.

**Figure. 4.**
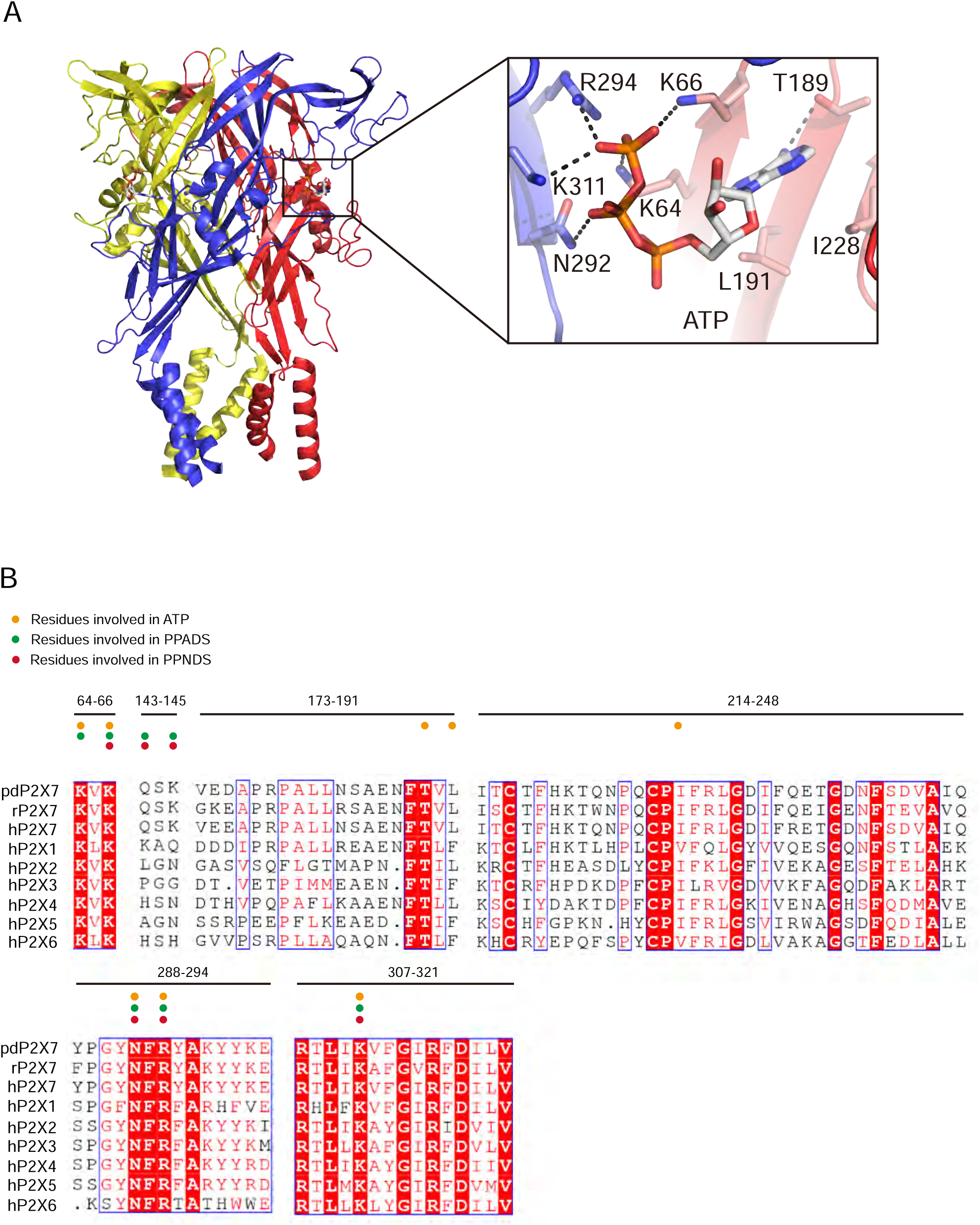
ATP binding site and sequence comparison. (A) Overall structure and close-up view of the ATP-bound rat P2X7 structure (PDB ID: 6U9W). The cytoplasmic domain is not shown. Dotted black lines indicate hydrogen bonding. (B) Sequence alignment of *Ailuropoda melanoleuca* P2X7 (pdP2X7) (Accession number: XP_002913164.3), *Rattus norvegicus* (rP2X7) (Accession number: Q64663.1) and *Homo sapiens* P2X receptors (P2X1: P51575.1, P2X2: Q9UBL9.1, P2X3: P56373.2, P2X4: Q99571.2, P2X5: Q93086.4, P2X6: O15547.2, and P2X7: Q99572.4). Orange, green and red circles indicate the residues involved in ATP, PPADS and PPNDS recognition.

In both structures, the pyridoxal phosphate group has extensive interactions with the receptor. In contrast, the other parts of the compounds, a naphthylazo group with two sulfonic acid groups and a nitro group (PPNDS) and an azophenyl group with two sulfonic acid groups (PPADS), have fewer interactions (**Fig. 2 and 3**).

The phosphate group of PPNDS and PPADS interacts directly with the side chain of Arg294 and possibly also with Lys145, possibly via a water molecule (**Fig. 2B and 3B and Fig. S7**). However, it should also be noted that it is difficult to conclude the existence of the water molecule at this site due to the limited resolution of our structures. Furthermore, in the PPADS-bound structure, Lys64 mediates an additional interaction with the phosphate group of Lys64. These extensive interactions between the phosphate group and the receptor resemble those with the phosphate groups of ATP (**Fig. 4A**). Furthermore, the Asn292 and Lys311 residues are similarly involved in the interaction with the hydroxyl group of the pyridoxal part of PPNDS and PPADS (**Fig. 2B and 3B**).

Interestingly, despite the structural differences between PPNDS and PPADS, the two common sulfonic acid groups form hydrogen bonds with the side chains of the same residues in the receptor, Lys66 and Gln143 (**Fig. 2B and 3B**).

Finally, to verify the binding mode of the pyridoxal phosphate derivatives, we performed molecular dynamics (MD) simulations of the higher-resolution PPNDS-bound structure embedded in lipids (**Fig. S8**). The overall structures were stable during the simulations, and PPNDS remained stably bound to the receptor (**Fig. S8**).

### Structural comparison and inhibition mechanism

To gain insights into the mechanisms of the orthosteric inhibition of P2X receptors by PPNDS and PPADS, we superimposed our structures and the previously determined ATP-bound P2X7 structure onto the apo-state P2X7 structure (**Fig. 5**). The PPNDS-bound structure and PPADS-bound structure are very similar, with 0.52 Å RMSD values for the Cα atoms of 960 residues. Only the comparison with the higher-resolution PPNDS-bound structure is described in the following discussion.

**Figure. 5.**
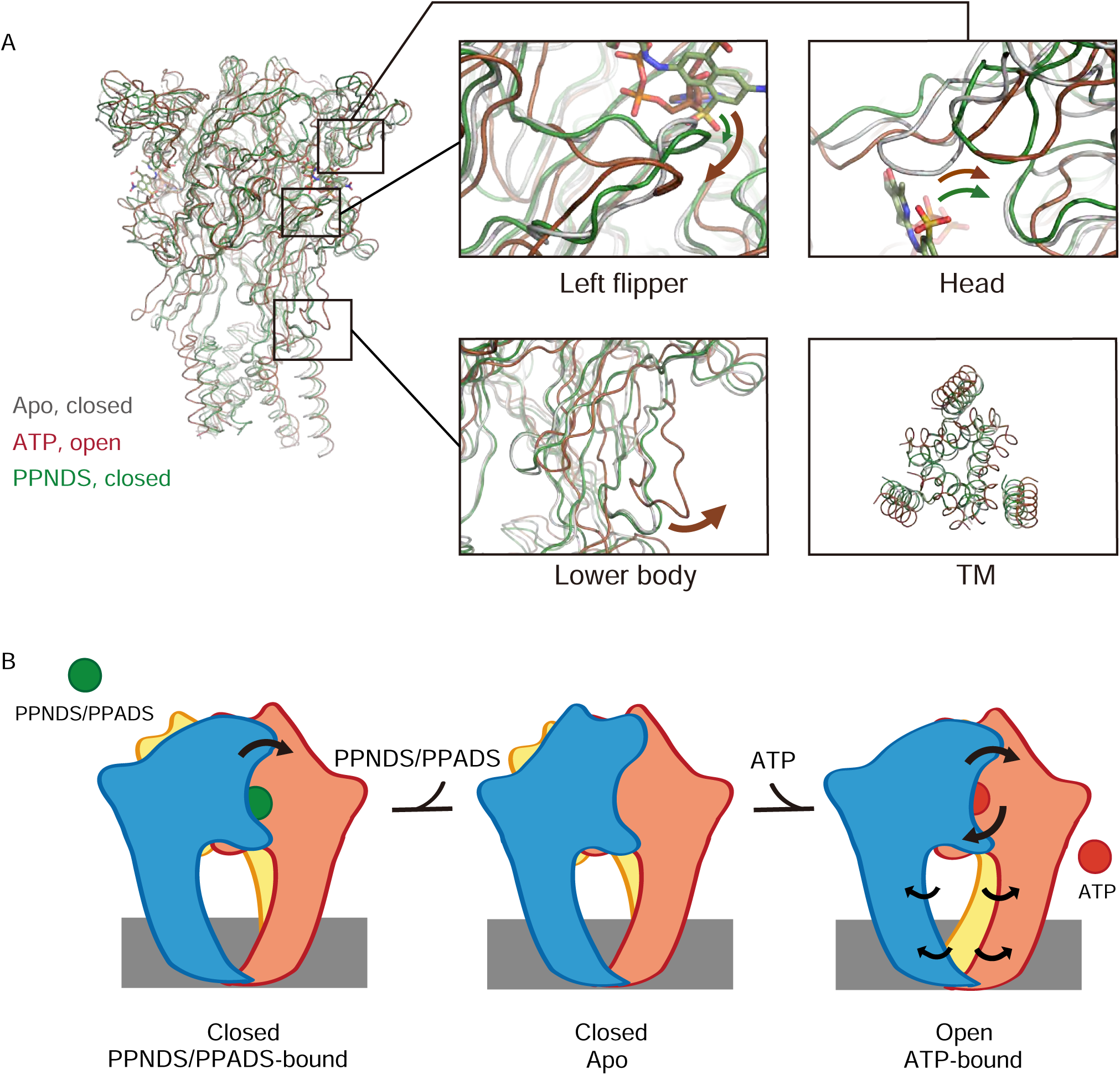
Structural comparison and inhibition mechanism. (A) Superposition of the ATP-bound rP2X7 structure (red, PDB ID: 6U9W) and the PPNDS-bound pdP2X7 structure (green, this study) onto the apo rP2X7 structure (gray, PDB ID: 6U9V). Close-up views of the head, left flipper, and lower body domains and the intracellular view of the TM domain are shown in each box. Arrows indicate the conformational changes from the apo to ATP-bound states (red) and from the apo to the PPNDS-bound states (green). (B) A cartoon model of the PPNDS/PPADS-dependent inhibition and ATP-dependent activation mechanisms.

First, the activation of P2X receptors from the apo (closed) state to the ATP-bound (open) state is known to require motions of both the head and left flipper domains^40, 41, 42^ (**Fig. 5A**). These motions are coupled to the movement of the lower body domain, which is directly connected to the TM domain for channel opening (**Fig. 5A**).

In the PPNDS-bound structure, while we observed motion of the head domain similar to that in the ATP-bound structure, there was only a small structural change in the left flipper domain (**Fig. 5A**). Consequently, there was no structural change in the lower body domain or associated gating motion of the TM domain (**Fig. 5A**). In the ATP-bound structure, the three phosphate groups of ATP in the U-shaped conformation pushed down the left flipper (**Fig. 4A and 5A**). In contrast, both PPNDS and PPADS possess only one phosphate group, so there was no corresponding downward movement in the left flipper domain (**Fig. 2 and 5A**).

To summarize, the structural comparison indicates how PPNDS and PPADS inhibit the ATP-dependent activation of P2X receptors (**Fig. 5B**). The binding of these competitive inhibitors may prevent the downward movement of the left flipper domain, which is required for channel opening (**Fig. 5B**).

### Structure-based mutational analysis

To analyze the mechanism of P2X receptor binding to the pyridoxal-5’-phosphate derivatives, we performed structure-based mutational analysis by whole-cell patch-clamp recording of pdP2X7 (**Fig. 6 and Fig. S1**). Most of the residues involved in PPNDS and PPADS binding overlap with the conserved residues involved in ATP binding (**Fig. 4B**). Thus, we did not generate mutants targeting these residues (Lys64, Lys66, Asn292, Arg294, and Lys311), as such mutations are known to severely affect or abolish ATP-dependent gating of P2X receptors^27, 37, 38, 39^. Instead, we aimed to mutate the residues surrounding the ATP-binding site, which differ among P2X receptor subtypes. Such residues may be important for the subtype-specific differences in the affinity of the pyridoxal-5’-phosphate derivative to P2X receptors. To design these mutants, we performed structural comparisons of our structures with the AlphaFold-based structural model of human P2X1^43^ (**Fig. 6A**), which has high affinity for both PPNDS and PPADS^21,35^. In particular, we compared the residues surrounding the ATP-binding site in our structure (Gln143, Val173, Ile214, Gln248 and Tyr288) with those in human P2X1 (hP2X1) (**Fig. 4B**). Based on these comparisons, we mutated these residues to the corresponding amino acid residues in hP2X1 (Gln143Lys, Val173Asp, Ile214Lys, Gln248Lys, and Tyr288Ser) or to alanine (Gln143Ala, Gln248Ala, and Tyr288Ala). Using these mutants, we performed whole-cell patch clamp recording to analyze the effect of PPNDS on these mutants (**Fig. 6B**).

**Figure. 6.**
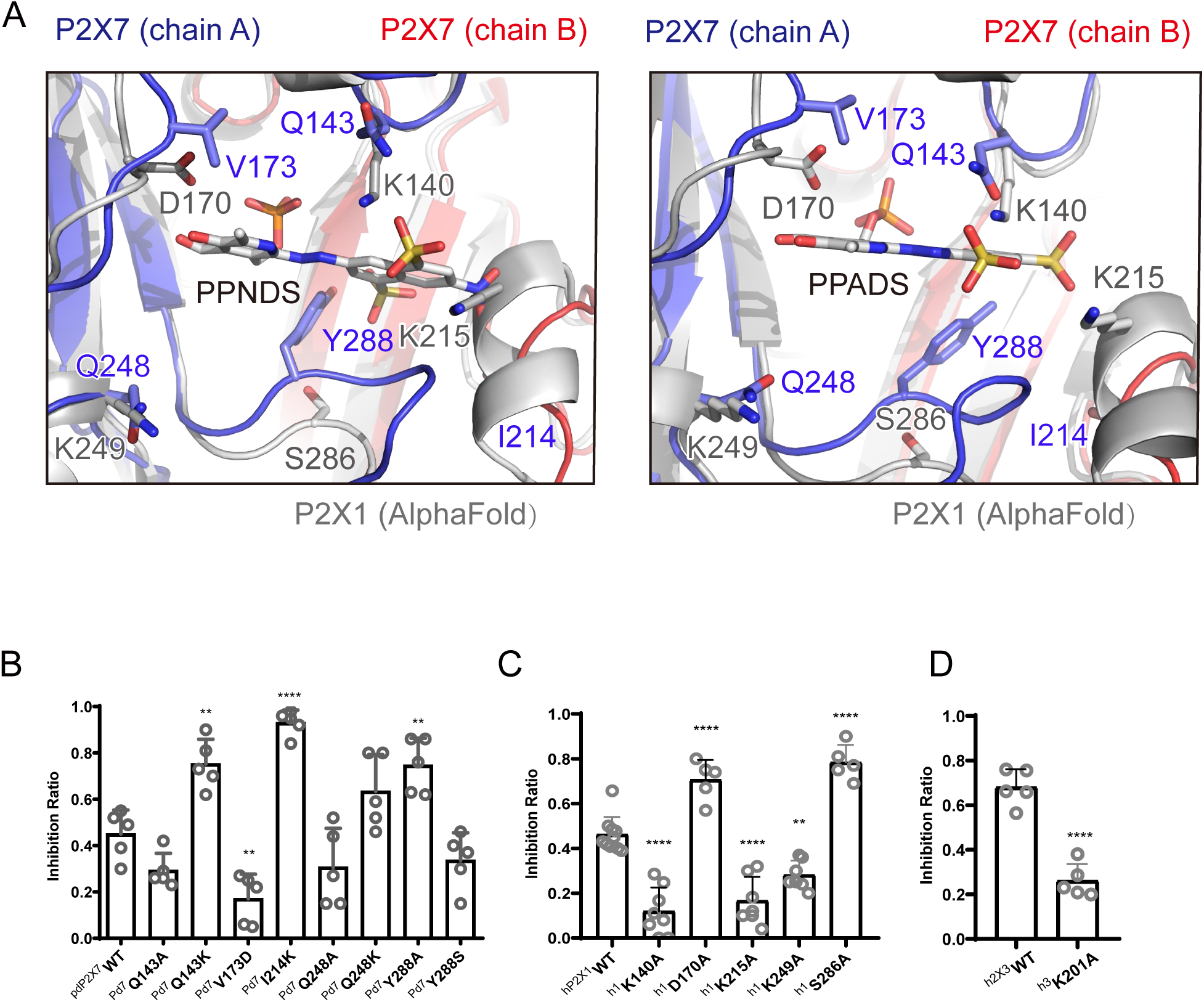
Structure-based mutational analysis. (A) Superimposition of the PPNDS-bound and PPADS-bound structures in this study onto the predicted human P2X1 structure (AlphaFold). Each subunit of the PPNDS-bound and PPADS-bound structures is shown in blue, yellow, and red, while the predicted human P2X1 structure is shown in gray. The PPNDS and PPADS molecules and the residues surrounding PPNDS and PPADS that are different between pdP2X7 and hP2X1 are shown as sticks. (B) Effects of PPNDS (10 µM) on ATP (1 mM)-evoked currents of pdP2X7 and its mutants (mean ± SD, n = 5). (C) Effects of PPNDS (1 µM) on ATP (1 µM)-evoked currents of hP2X1 and its mutants (mean ± SD, n = 5-10). (D) Effects of PPNDS (10 µM) on ATP (1 µM)-evoked currents of hP2X3 and its mutants (mean ± SD, n = 5, one-way ANOVA post hoc test, **: p <0.01, ****: p <0.0001 vs. WT.).

In the mutational analysis, among three mutations to the lysine residue, two (Gln143Lys and Ile214Lys) showed significantly increased sensitivity to PPNDS (**Fig. 6B**). In addition, the Gln248Lys mutant showed slightly higher sensitivity to PPNDS, but not as significant an increase as the other two mutants (**Fig. 6B**). Interestingly, while Tyr288 in pdP2X7 corresponds to Ser286 in hP2X1, the mutation of Tyr288 to alanine instead of serine significantly increased the affinity for PPNDS (**Fig. 6B**). In addition, mutating Val173 in pdP2X7 to aspartate significantly reduced the sensitivity to PPNDS (**Fig. 6B**). The two mutants Gln143Ala and Gln248Ala showed little or no decrease in PPNDS sensitivity (**Fig. 6B**).

Following the result of the mutation analysis of pdP2X7, we then designed alanine mutants of hP2X1 at Lys140 (Gln143 in pdP2X7), Asp170 (Val173 in pdP2X7), Lys215 (Ile214 in pdP2X7), Lys249 (Gln248 in pdP2X7), and Ser286 (Tyr288 in pdP2X7) (**Fig. 4B and 6A**). Using these mutants, we then performed whole-cell patch-clamp recording to evaluate the effect of PPNDS on these mutants (**Fig. 6C**). All three mutants at the lysine residues (Lys140Ala, Lys215Ala and Lys249Ala), especially the Lys140Ala and Lys215Ala mutants, showed a significant decrease in PPNDS sensitivity (**Fig. 6C**). This result is largely consistent with the corresponding lysine-substituted mutants of pdP2X7 (**Fig. 6B**). Interestingly, the mutation of Asp170 and Ser286 to alanine increased the sensitivity to PPNDS (**Fig. 6C**). Consistently, in the mutational analysis of the corresponding residues in pdP2X7, the mutation of Val173 to aspartate decreased PPNDS sensitivity, and the mutation of Tyr288 to alanine increased PPNDS sensitivity (**Fig. 6B**).

Finally, among the three Lys residues involved in PPNDS sensitivity in hP2X1 (Lys140, Lys215, Lys249), Lys215 is also conserved in hP2X3 (Lys201). Accordingly, we performed mutational analysis of the hP2X3 Lys201Ala mutant (**Fig. 6D**). As expected, the Lys201 mutant showed a decrease in PPNDS sensitivity (**Fig. 6D**).

In summary, our mutational analysis based on structural comparison and sequence alignment identified several key residues involved in PPNDS sensitivity, particularly the residues involved in the subtype-specific difference in affinity for PPNDS.

## Discussion

In this work, we determined the cryo-EM structure of pdP2X7 in complex with two classical non-ATP analog inhibitors, PPNDS and PPADS, of pyridoxal phosphate derivatives (**Fig. 1**) and revealed their orthosteric binding site (**Fig. 2 and 3**). The binding site for PPNDS and PPADS has high overlap with the ATP binding site (**Fig. 4**). In the cryo-EM structures, the phosphate group of PPNDS and PPADS appears to occupy the position of the γ-phosphate group of ATP in the ATP-bound structure (**Figs. 2-4**). Comparison with the previously reported apo (closed) and ATP-bound (open) structures showed that, in contrast to ATP binding, the binding of PPNDS and PPADS does not induce the downward motion of the left flipper, which is essential for channel activation (**Fig. 5**). These observations provide mechanistic insights into channel inhibition by pyridoxal phosphate derivatives. Finally, structure-based mutational analyses revealed several key residues important for PPNDS sensitivity, particularly for the subtype-specific difference in sensitivity (**Fig. 6**).

Besides PPADS and PPNDS in this study, TNP-ATP is well known as a classical competitive inhibitor for P2X receptors^21^. In the previously reported TNP-ATP bound P2X7 structure, TNP-ATP binding induces the conformational changes of the left flipper region but not of the head domain^31^. In contrast, PPADS and PPNDS binding induces the conformational changes of the head domain but not of the left flipper region (**Fig. 5**). These contrasts would highlight the uniqueness of the competitive inhibition mechanism by pyridoxal phosphate derivatives as well as the diversity of competitive inhibition mechanisms of P2X receptors.

There are several previous reports characterizing the P2X receptor binding site for pyridoxal phosphate derivatives, especially PPADS^15, 16, 18, 19^. The mutation of Glu249 in rat P2X4 (rP2X4) to lysine was designed based on the corresponding residue in hP2X1 (Lys249) to confer PPADS sensitivity on the PPADS-insensitive P2X4 receptor^18^. Glu249 in rP2X4 corresponds to Gln248 in pdP2X7 (**Fig. 4B**). In our structures, this residue is proximal to the hydroxyl moieties of the pyridoxal phosphate group of PPADS and PPNDS (**Fig. 6A**). More recently, the combination of docking simulation and electrophysiology showed that Lys70, Asp170, Lys190 and Lys249 participate in PPADS binding in hP2X1^15^. Among these four residues, Lys70 and Lys190 are directly involved in ATP binding^27^, and Lys249 corresponds to Glu249 in rP2X4, as shown in a previous study^18^. In addition, the mutation of Asp170 to cysteine increased sensitivity to PPADS^15^. Consistently, our mutational analysis revealed that the Asp170Ala mutation in hP2X1 also increased the sensitivity to PPNDS (**Fig. 6C**). According to our structures, the corresponding residue in pdP2X7 (Val173) is located proximal to the pyridoxal phosphate group of PPADS, and the pyridoxal phosphate group is in common with that of PPADS (**Fig. 6A**). Therefore, Asp170 in hP2X1 is involved in sensitivity to both PPADS and PPNDS.

It should be noted that the orientation of PPADS in the recent docking model is very different from that in our cryo-EM structure. In the docking model, compared to our cryo-EM structure, PPADS shows an almost 180-degree rotation relative to the axis of rotation parallel to the membrane with the azide group of PPADS as the fulcrum^15^. This difference in the orientation of the compound may explain why hP2X1 Lys140 and Lys215 were not predicted to be involved in PPADS binding in the previous docking model, in contrast to the results of our study.

In the past, several types of pyridoxal phosphate derivatives have been identified as P2X inhibitors^13, 23^. However, most of them share a common weakness in subtype specificity, where they tend to show high affinity for both P2X1 and P2X3. For example, MRS 2257 was identified as the most active PPADS analog from the screening but has IC50 values of 5 nM and 22 nM for P2X1 and P2X3, respectively^22^. In addition, another PPADS analog, termed 36j, shows improved subtype specificity for P2X3 (IC50: 60 nM for P2X3) but still shows a moderate inhibitory effect on P2X1 at 10 µM^23^. Thus, it is still difficult to obtain competitive inhibitors of P2X receptors, including pyridoxal phosphate derivatives, with subtype specificity. This is probably because the residues directly involved in ATP binding are strictly conserved among the subtypes (**Fig. 4B**). To overcome this situation, our work might facilitate the rational design of pyridoxal phosphate derivatives with strict subtype specificity for P2X receptors. We have not only defined the binding mode of pyridoxal phosphate derivatives to P2X receptors but also newly identified a subtype-specific residue for pyridoxal phosphate derivatives. For example, Lys215 and Lys249 in hP2X1 are important for PPNDS sensitivity (**Fig. 6C**) but are not conserved in other P2X subtypes, including P2X3 (**Fig. 4B**). These findings would provide a clue for the design of more subtype-specific pyridoxal phosphate derivatives.

In conclusion, our structural and functional analyses provided mechanistic insights into the orthosteric inhibition mechanism of P2X receptors by the classical pyridoxal phosphate derivative P2X antagonist. In addition, we identified key residues involved in compound sensitivity, especially the differential sensitivity between P2X subtypes. These results potentially facilitate the development of subtype-specific compounds targeting P2X receptors, which have attracted widespread interest as therapeutic targets.

## Methods

### Expression and purification of P2X7

The previously reported functional expression construct of giant panda (*Ailuropoda melanoleuca*) P2X7 for structural studies (pdP2X7, residues 22-359, N241S/N284S/V35A/R125A/E174K, XP_002913164.1)^29^ was synthesized (Genewiz, China), subcloned, and inserted into a modified version of the pFastBac vector (Invitrogen, USA) with an octahistidine tag, Twin-Strep-tag, mEGFP, and a human rhinovirus (HRV) 3C protease cleavage site at the N-terminus. Using the Bac-to-Bac system, the mEGFP-fusion pdP2X7 construct was expressed in Sf9 cells infected with baculovirus. The Sf9 cells were collected by centrifugation (5,400 × g, 10 min) and subsequently disrupted using an ultrasonic homogenizer in TBS buffer (20 mM Tris pH 8.0, 150 mM NaCl) containing 1 mM phenylmethylsulfonyl fluoride (PMSF), 5.2 μg/mL aprotinin, 1.4 μg/mL pepstatin, and 1.4 μg/mL leupeptin. The supernatant was harvested after centrifugation (7,600 × g, 20 min). The membrane fraction was then isolated by ultracentrifugation (200,000 × g, 1 h) and solubilized in buffer A (50 mM Tris pH 7.5, 150 mM NaCl) containing 2% (w/v) n-dodecyl-beta-D-maltopyranoside (DDM) at 4 °C for 1 hour. The solubilized supernatant was collected by another round of ultracentrifugation (200,000 × g, 1 h) and applied to a Strep-Tactin resin column (Qiagen, USA) equilibrated with buffer A containing 0.025% (w/v) DDM. The resin was incubated for 1 hour, and the column was eluted with buffer B (100 mM Tris pH 8.0, 150 mM NaCl, 2.5 mM desthiobiotin, 0.025% (w/v) DDM). The eluted protein was concentrated to 1 mg/ml before being prepared for nanodisc reconstitution.

### Nanodisc reconstitution

Soybean polar lipid extract (Avanti Polar Lipids, USA) was dissolved in chloroform, dried under a nitrogen stream, and then resuspended in reconstitution buffer (20 mg/ml soybean polar lipid, 20 mM HEPES pH 7.0, and 150 mM NaCl). Following a 1-hour incubation at room temperature, the lipid suspension was subjected to sonication for 5 minutes until the lipids reached a near-transparent state. Subsequently, DDM (Anatrace, USA) was added at a final concentration of 0.4% and incubated at room temperature overnight. The mEGFP-fusion pdP2X7, MSP2N2 protein, and soybean polar lipid were combined in a molar ratio of 1:3:180. This mixture was then incubated at 4 °C for 1 hour and further subjected to a 4-hour incubation with bio-beads (Bio-Rad, USA). After incubation, the bio-beads were removed via filtration, and the nanodisc fractions containing mEGFP-fusion pdP2X7 were bound to Ni-NTA (Qiagen, USA) resin preequilibrated with wash buffer (20 mM HEPES pH 7.5, 150 mM NaCl, 30 mM imidazole) and subsequently eluted using elution buffer (20 mM HEPES pH 7.5, 150 mM NaCl, 300 mM imidazole). To cleave the N-terminal EGFP, the elution was mixed with HRV3C protease and incubated at room temperature for 1 hour, followed by overnight incubation at 4 °C. The nanodisc-reconstituted pdP2X7 protein was separated through size-exclusion chromatography using a Superdex 200 Increase 10/300 column (Cytiva, USA) preequilibrated with SEC buffer (20 mM HEPES pH 7.5 and 150 mM NaCl) and subsequently concentrated to 0.9 mg/ml. P2X7 antagonists (PPNDS or PPADS) were added to the nanodisc-reconstituted pdP2X7 at a final concentration of 50 μM and incubated on ice for 1 hour before cryo-EM grid preparation.

### EM data acquisition

For both the PPNDS-bound and PPADS-bound pdP2X7 samples, a total of 2.5 μl of the nanodisc-reconstituted pdP2X7 was applied to a glow-discharged holey carbon-film grid (Quantifoil, Au 1.2/1.3, 300 mesh, USA). The grid was then blotted using a Vitrobot (Thermo Fisher Scientific, USA) system with a 3.0-second blotting time at 100% humidity and 4 °C, followed by plunge-freezing in liquid ethane. Cryo-EM data collection was carried out using a 300 kV Titan Krios microscope (Thermo Fisher Scientific, USA) equipped with a K3 direct electron detector (Gatan Inc., USA). The specimen stage temperature was maintained at 80 K. Movies were recorded using beam-image shift data collection methods^44^ in superresolution mode, with a pixel size of 0.41 Å (physical pixel size of 0.83 Å), a magnification of 29,000, and defocus values ranging from -1.3 µm to - 2.0 µm. The dose rate was set to 20 e-s^−1^, and each movie consisted of 40 frames with an exposure of 50 e-Å^−2^, resulting in each movie being 1.724 s long.

### Image processing

A total of 9,664 and 4,692 movies for the PPNDS-bound and PPADS-bound pdP2X7 samples, respectively, were motion-corrected and binned with MotionCor2^45^ with a patch of 5 × 5, producing summed and dose-weighted micrographs with a pixel size of 0.83 Å. Contrast transfer function (CTF) parameters were estimated by CTFFIND 4.1^46^. Particle picking and 2D classification were performed using RELION 3.1^47^. In total, 1,537,753 particles for the PPNDS-bound sample and 1,835,907 particles for the PPADS-bound sample were autopicked and extracted using a box size of 256 × 256 pixels. After 2D classification, we performed 3D classification with C1 symmetry using RELION 3.1 on 633,674 particles for the PPNDS-bound sample and 236,753 particles for the PPADS-bound sample. Then, 121,008 particles for the PPNDS-bound sample and 161,188 particles for the PPADS-bound sample were selected for nonuniform refinement using cryoSPARCv4.2.1^48^, applying C3 symmetry for the final 3D reconstruction. The resulting resolutions of the PPNDS-bound and PPADS-bound pdP2X7 structures were 3.3 Å and 3.6 Å, respectively, as determined by the Fourier shell correlation (FSC) = 0.143 criterion on the corrected FSC curves. The local resolution was estimated using cryoSPARCv4.2.1. The workflows for image processing and for 3D reconstruction are shown in **Figs. S2-S5**. The figures were generated by UCSF Chimera^49^.

### Model building

The initial models of pdP2X7 were manually built starting from the previously reported pdP2X7 structure (PDB ID: 5U1L). Manual model building was performed using Coot^50^. Real-space refinement was performed using PHENIX^51^. All structure figures were generated using PyMOL (https://pymol.org/). For the predicted structure of human P2X1, the previously generated model using AlphaFold and ColabFold was used^43, 52, 53^. The sequence alignment figure was generated using Clustal Omega^54^ and ESPript 3.0^55^.

### Electrophysiology

Human embryonic kidney 293 (HEK293) cells were purchased from Shanghai Institutes for Biological Sciences and cultured in Dulbecco’s modified Eagle’s medium supplemented with 10% fetal bovine serum (FBS), 1% penicillin‒streptomycin, and 1% GlutaMAX™ at 37 °C in a humidified atmosphere of 5% CO_2_ and 95% air^56,57^. Plasmids harboring hP2X1, hP2X3 or pdP2X7 were transfected into cells by calcium phosphate transfection^58^. Currents of hP2X1 and hP2X3 were recorded using nystatin (Sangon Biotech, China) perforated recordings to prevent rundown in current during multiple dose applications of ATP. Nystatin (0.15 mg/mL) was diluted with a high-potassium internal intracellular solution containing 75 mM K_2_SO_4_, 55 mM KCl_2_, 5 mM MgSO_4_, and 10 mM HEPES (pH 7.4). Currents of PdP2X7 receptors were recorded using a conventional whole-cell patch configuration. After 24–48 h of transfection, HEK293 cells were recorded at room temperature (25 ± 2 °C) using an Axopatch 200B amplifier (Molecular Devices, USA) with a holding potential of -60 mV. Current data were sampled at 10 kHz, filtered at 2 kHz, and analyzed using PCLAMP 10 (Molecular Devices, USA) for analysis. HEK293 cells were bathed in standard extracellular solution (SS) containing 2 mM CaCl_2_, 1 mM MgCl_2_, 10 mM HEPES, 150 mM NaCl, 5 mM KCl, and 10 mM glucose with the pH adjusted to 7.4. For conventional whole-cell recordings, the pipette solutions consisted of 120 mM KCl, 30 mM NaCl, 0.5 mM CaCl_2_, 1 mM MgCl_2_, 10 mM HEPES, and 5 mM EGTA with pH adjusted to 7.4. ATP and other compounds were dissolved in SS for P2X1 and P2X3 and applied to Y-tubes. For pdP2X7, ATP and other compounds were dissolved in 0Ca, 0Mg solution containing 150 mM NaCl, 10 mM glucose, 10 mM HEPES, 5 mM KCl, and 10 mM EGTA with the pH adjusted to 7.4^59^. PPNDS was purchased from APE ×BIO, and PPADS was purchased from MCE. The standard solution and 0Ca, 0Mg solution were formulated with compounds from Aladdin, and internal solutions were formulated with compounds from Sigma‒Aldrich^60^. All electrophysiological recordings were analyzed using Clampfit 10.6 (Molecular Devices, USA). Pooled data are expressed as the mean and standard error (s.e.m.). Statistical comparisons were made using Bonferroni’s post hoc test (ANOVA). ∗∗ p < 0.01 and ∗∗∗∗ p < 0.0001 were considered significant.

### Molecular dynamics simulations

The energy-minimized models of the PPNDS-bound pdP2X7 were used as the initial structures for MD simulations. A large 1-palmitoyl-2-oleoyl-sn-glycero-3-phosphocholine (POPC, 300 K) bilayer, available in System Builder of DESMOND^58,61^, was built to generate a suitable membrane system based on the OPM database^62^. The systems were dissolved in simple point charge (SPC) water molecules. The DESMOND default relaxation protocol was applied to each system prior to the simulation run. 1) 100 ps simulations in the NVT (constant number (N), volume (V), and temperature (T)) ensemble with Brownian kinetics using a temperature of 10 K with solute heavy atoms constrained; 2) 12 ps simulations in the NVT ensemble using a Berendsen thermostat with a temperature of 10 K and small-time steps with solute heavy atoms constrained; 3) 12 ps simulations in the NPT (constant number (N), pressure (P), and temperature (T)) ensemble using a Berendsen thermostat and barostat for 12 ps simulations at 10 K and 1 atm, with solute heavy atoms constrained; 4) 12 ps simulations in the NPT ensemble using a Berendsen thermostat and barostat at 300 K and 1 atm with solute heavy atoms constrained; and 5) 24 ps simulations in the NPT ensemble using a Berendsen thermostat and barostat at 300 K and 1 atm without constraint. After equilibration, the MD simulations were performed for 0.3 µs. The long-range electrostatic interactions were calculated using the smooth particle grid Ewald method. The trajectory recording interval was set to 200 ps, and the other default parameters of DESMOND were used in the MD simulation runs. All simulations used the all-atom OPLS_2005 force field^63–65^, which is used for proteins, ions, lipids and SPC waters. The Simulation Interaction Diagram (SID) module in DESMOND was used for exploring the interaction analysis between PPNDS and pdP2X7. All simulations were performed on a DELL T7920 with NVIDIA TESLTA K40C or CAOWEI 4028GR with NVIDIA TESLTA K80. The simulation system was prepared, and the trajectory was analyzed and visualized on a CORE DELL T7500 graphics workstation with 12 CPUs.

### Statistics

Electrophysiological recordings were repeated 5-10 times. Error bars represent the standard error of the mean. Cryo-EM data collection and refinement statistics are summarized in **Table S1.**

## Data availability

The atomic coordinates and structural factors for the pdP2X7 in complex with PPNDS (PDB: 8JV8 and EMD-36671) and PPADS (PDB: 8JV7 and EMD-36670) have been deposited in the Protein Data Bank. All other relevant data are included in the paper or its supplementary material files, including the supplementary data file (**Data S1**), or deposited in ScienceDB (doi:10.57760/sciencedb.11168).

## Acknowledgments

We thank the staff scientists at the Center for Biological Imaging, Institute of Biophysics, Chinese Academy of Sciences for technical assistance with cryo-EM data collection (project numbers: CBIapp202007004; 2020-NFPS-PT-005280). This work was supported by funding from the National Natural Science Foundation of China to M.H. (32071234, 32271244 and 32250610205). This work was also supported by the Innovative Research Team of High-level Local Universities in Shanghai, a key laboratory program of the Education Commission of Shanghai Municipality (ZDSYS14005). This work was also supported by JST, PRESTO Grant Number JPMJPR20E1, Japan to M.I.

## Author contributions

D.S. expressed and purified P2X7 and performed cryo-EM experiments with assistance from F.J., Y.W., and M.I. D.S. and M.H. performed model building. C.Y. performed the electrophysiology experiments and MD simulation with assistance from Y.Y. and C.G. D.S., C.Y., C.G. and M.H. wrote the manuscript. C.G. and M.H. supervised the research. All authors discussed the manuscript.

