## Supplementary material for "Structural insights into the orthosteric inhibition of P2X receptors by non-ATP-analog antagonists": Figs. S1-8 and Table S1.

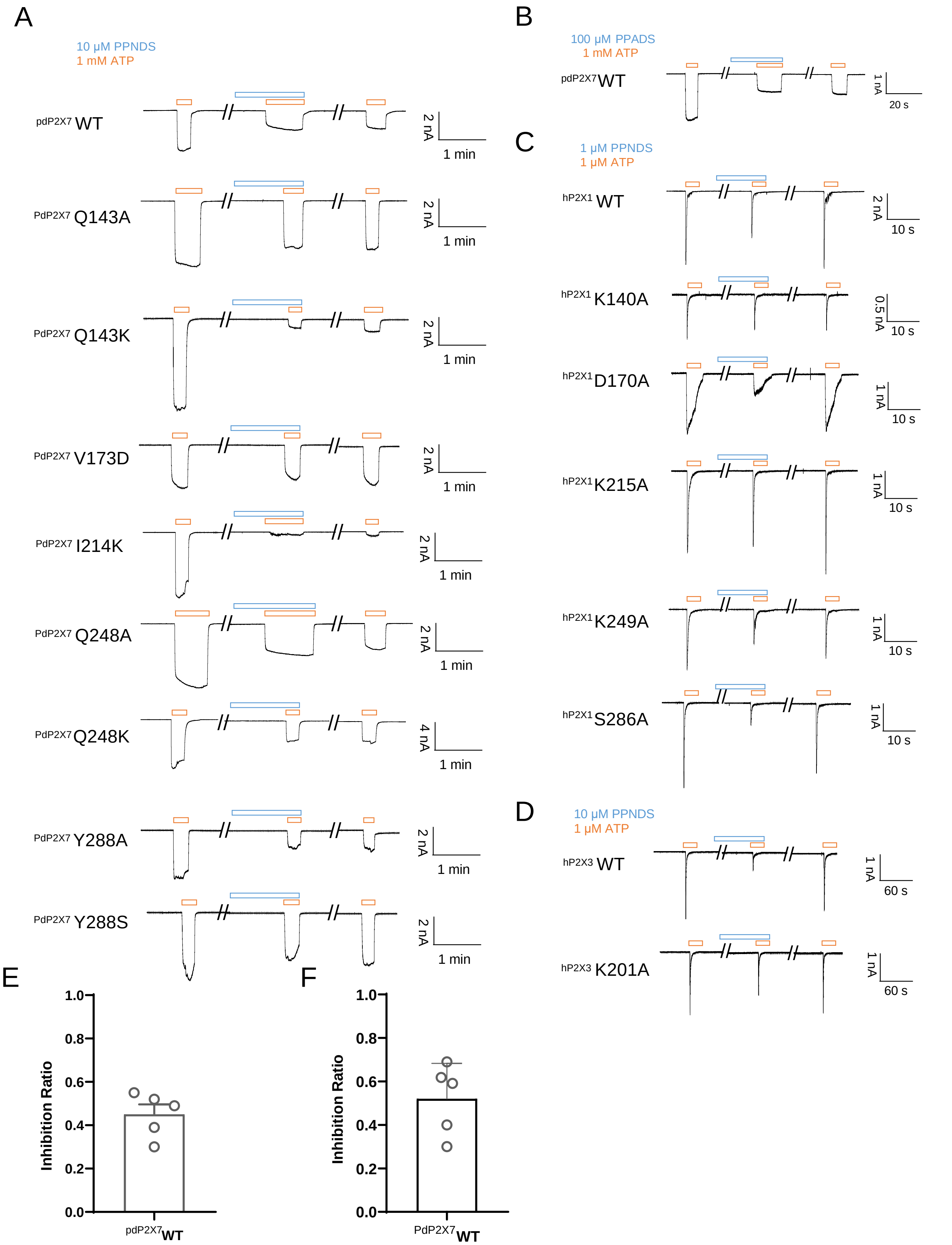


**Fig. S1. Effects of PPNDS and PPADS on P2X receptors by patch clamp recording**
(A-D) Representative current traces from patch clamp recordings of P2X receptors. Effects of 10 μM PPNDS (blue) on the 1 mM ATP-evoked (orange) current of pdP2X7 and its mutants (A). Effects of 100 μM PPADS (blue) on the 1 mM ATP-evoked (orange) current of pdP2X7 (B). Effects of 1 μM PPNDS (blue) on the 1 μM ATP-evoked (orange) current of hP2X1 (C). Effects of 10 μM PPNDS (blue) on the 1 μM ATP-evoked (orange) current of hP2X3 (D). (E-F) Effects of PPNDS (10 μM) (E) and PPADS (100 μM) (F) on ATP (1 mM)-evoked currents of pdP2X7 (mean ± SD, n = 5). The graph for PPNDS was taken from Fig. 6B. The inhibition ratio is defined by normalizing the peak current amplitude from the coapplication of PPNDS/PPADS and ATP to the peak current amplitude from the ATP application prior to the coapplication of PPNDS/PPADS and ATP.

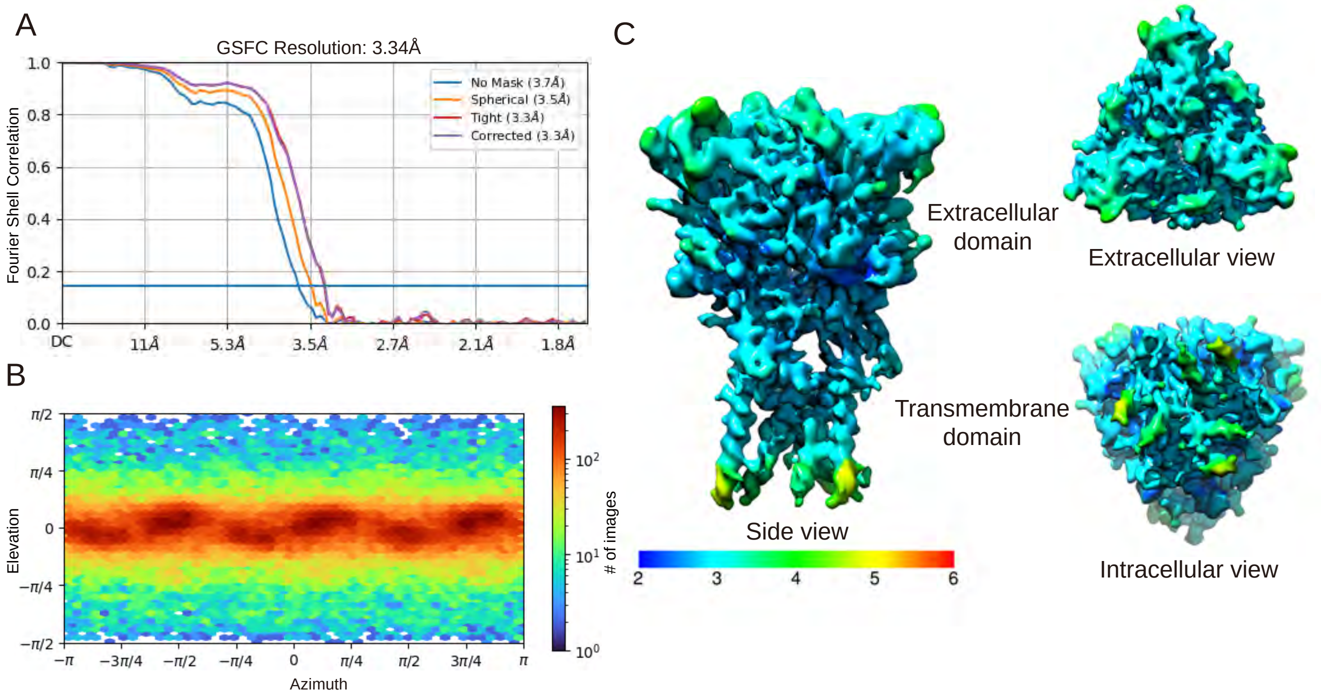


**Fig. S2. Cryo-EM analysis of PPNDS-bound pdP2X7**

(A) The gold-standard Fourier shell correlation curves for the PPNDS -bound data. (B) Angular particle distribution. The heat map of particle projections in each viewing angle. (C) The side view, a top-down view from the extracellular surface and a bottom-up view from the intracellular surface colored by local resolution.

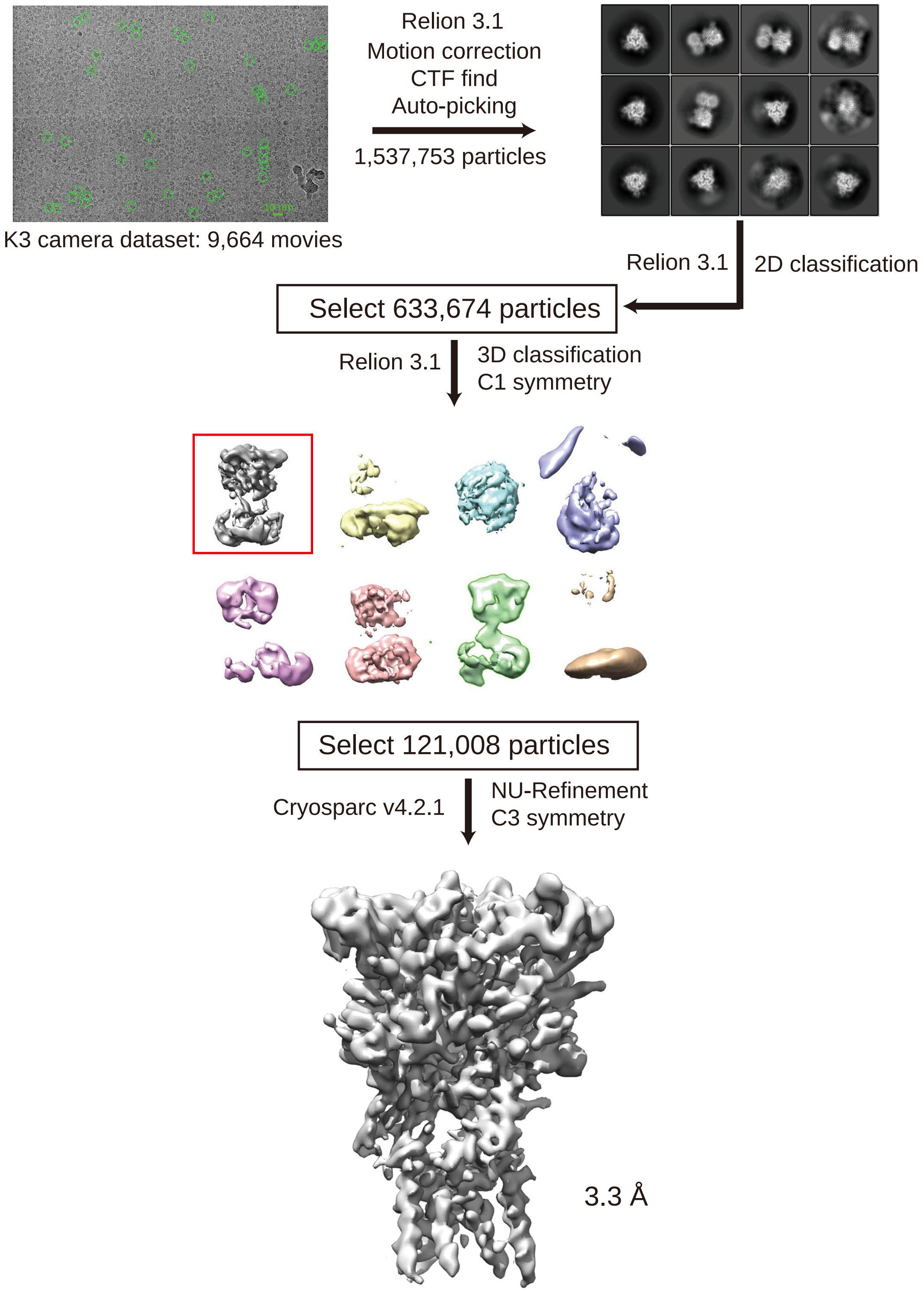


**Fig. S3. Cryo-EM data process for PPNDS-bound pdP2X7**

Before applying C3 symmetry, all steps including 3D classification were performed in Relion 3.1. With C3 symmetry, further refinement using Cryosparc v4.2.1 by non-uniform refinement of this final set of particles resulted in a cryo-EM map at 3.34 Å resolution.

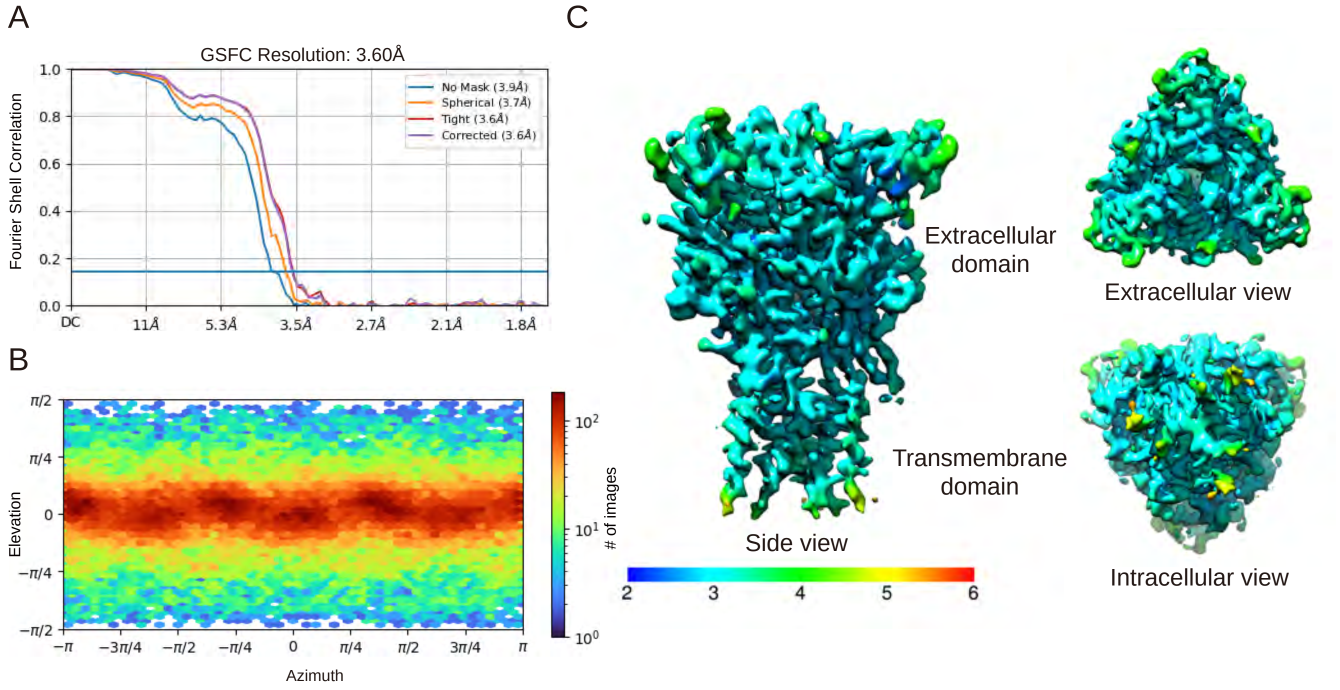


**Fig. S4. Cryo-EM analysis of PPADS-bound pdP2X7**

(A) The gold-standard Fourier shell correlation curves for the PPADS-bound data. (B) Angular particle distribution. The heat map of particle projections in each viewing angle. (C) The side view, a top-down view from the extracellular surface and a bottom-up view from the intracellular surface colored by local resolution.

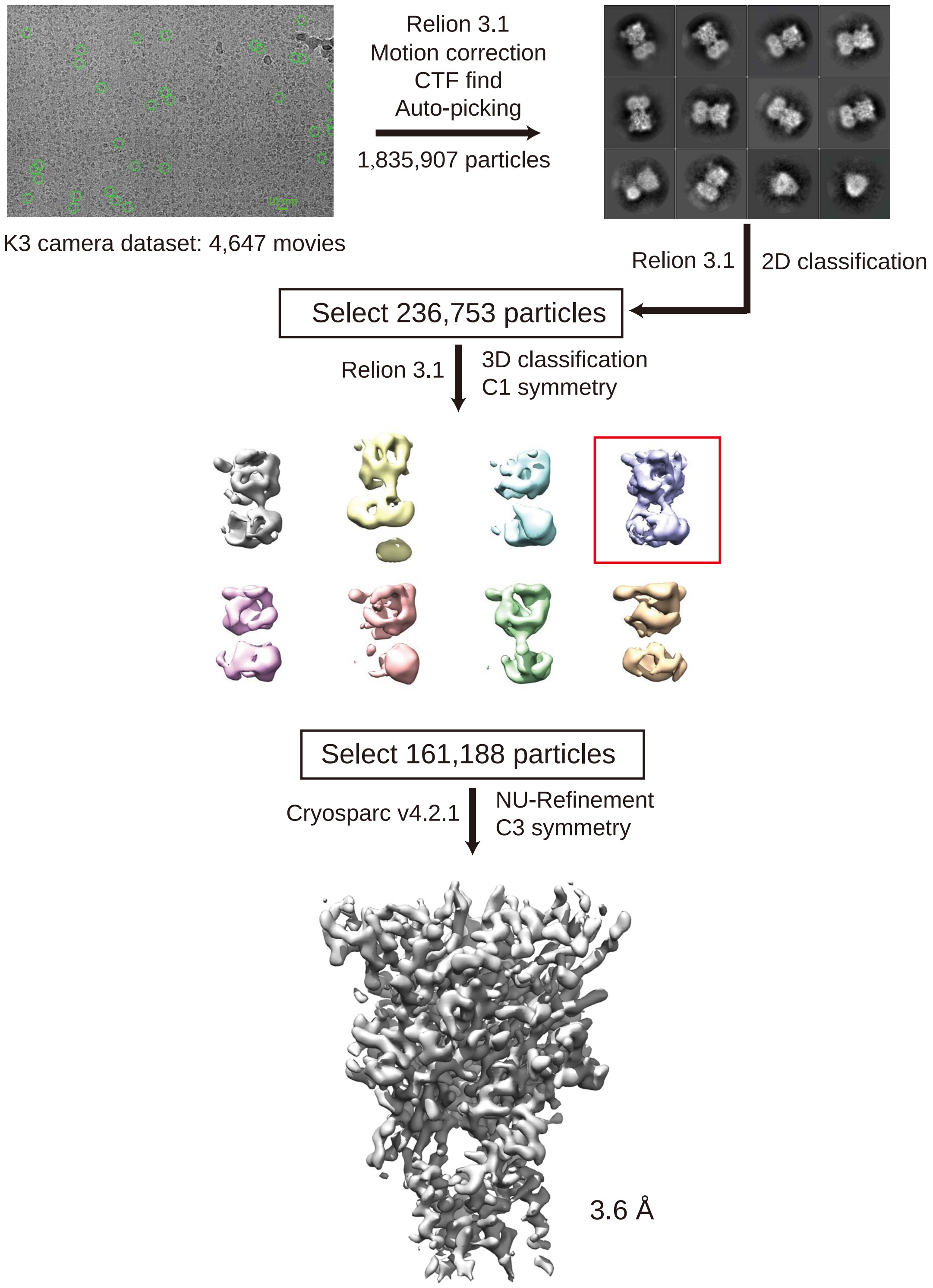


**Fig. S5. Cryo-EM data process for PPADS-bound pdP2X7**
Before applying C3 symmetry, all steps including 3D classification were performed in Relion 3.1. With C3 symmetry, further refinement using Cryosparc v4.2.1 by non -uniform refinement of this final set of particles resulted in a cryo-EM map at 3.60 Å resolution.

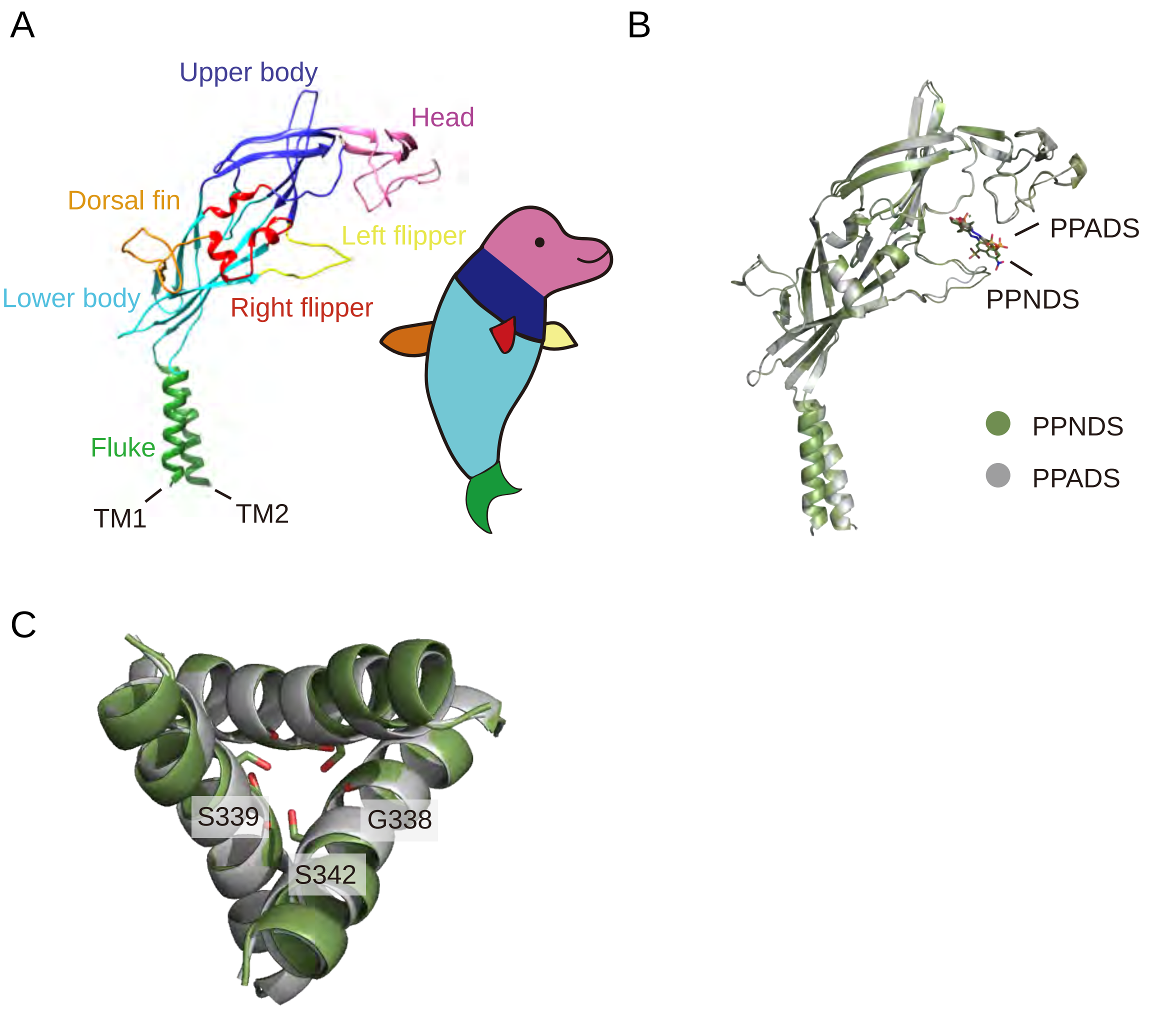


**Fig. S6. Dolphin model**

(A) The P2X7 protomer in cartoon representation. Each structural feature is colored according to the dolphin model. (B , C) Superposition of the PPNDS -bound structure (green) onto the PPADS -bound structure (gray). Each protomer is shown in cartoon representations, and PPNDS and PPADS are shown in stick representations (B). The intracellular view of thetransmembrane domain and the residues at the constriction regio are shown in stick representations (C).

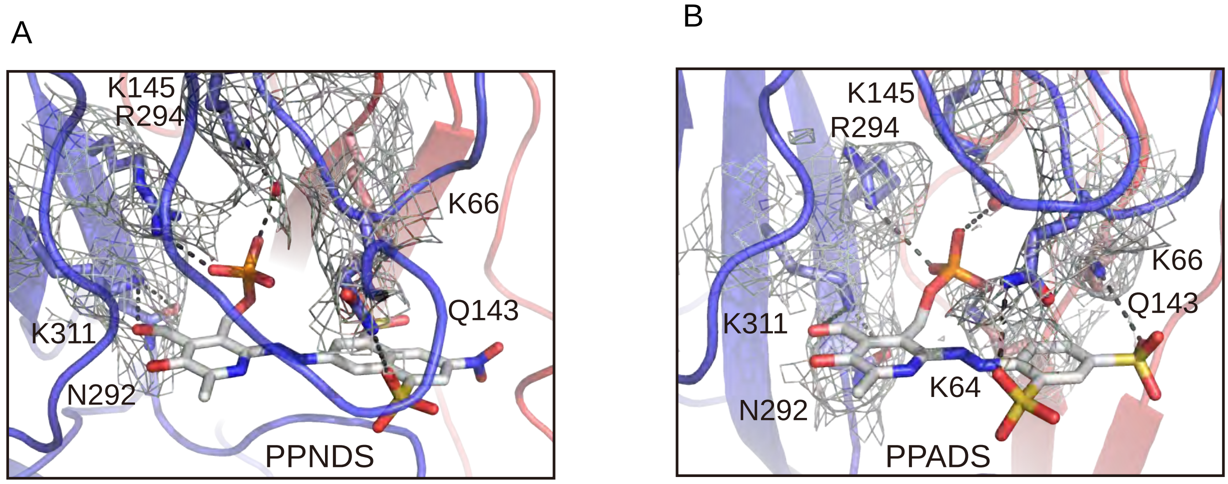


**Fig. S7. EM density maps for the PPNDS and PPADS binding sites**

(A, B) Close -up views of the PPNDS (A) and PPADS (B) binding sites. Dotted lines represent hydrogen bonds. The EM density maps for the residues involved in the PPNDS and PPADS interactions are shown and contoured at 5.0 σ and 4.0 σ, respectively.

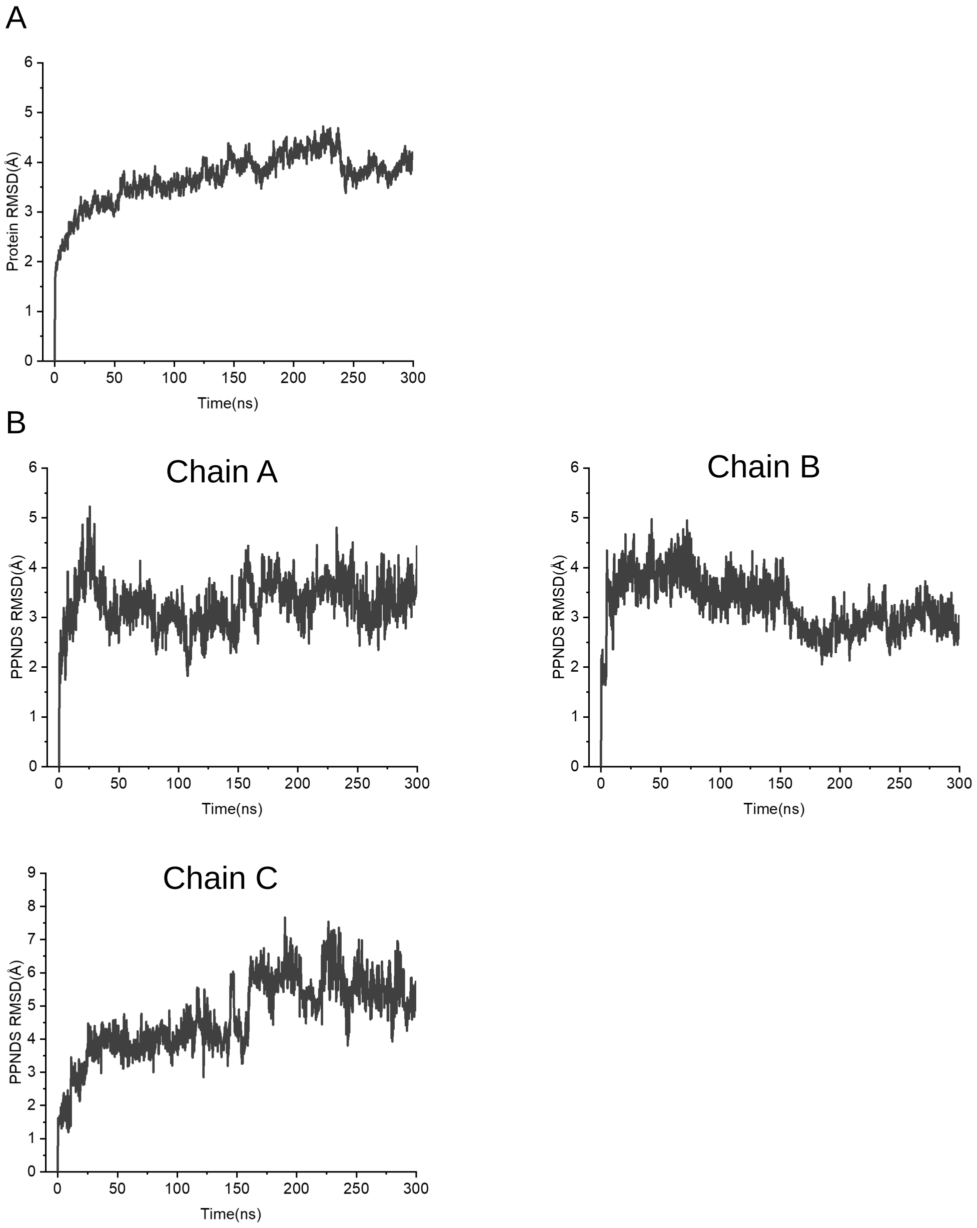


**Fig. S8. MD simulations of the PPNDS-bound pdP2X7 structure**
(A, B) The plots of the root mean square deviations (RMSD) of Cα atoms (A) and the RMSD values of atoms in PPNDS (B).

**Table S1. Cryo-EM data collection, refinement and validation statistics**

| | PdP2X7 w. PPNDS (EMD-36671) (PDB: 8JV8) | PdP2X7 w. PPADS (EMD-36670) (PDB: 8JV7) |
|---|---|---|
| **Data collection and processing** | | |
| Magnification | 29,000x | 29,000x |
| Voltage (kV) | 300 | 300 |
| Electron exposure (e–/Å$^2$) | 50 | 50 |
| Defocus range (μm) | -1.3 to -2.0 | -1.3 to -2.0 |
| Pixel size (Å) | 0.83 | 0.83 |
| Symmetry imposed | C3 | C3 |
| Initial particle images (no.) | 663,674 | 236,753 |
| Final particle images (no.) | 121,008 | 161,188 |
| Map resolution (Å) | 3.34 | 3.60 |
| FSC threshold | 0.143 | 0.143 |
| Map resolution range (Å) | 1.9-40.4 | 2.2-10.2 |
| **Refinement** | | |
| Initial model used (PDB code) | 5U1L | 5U1L |
| Model resolution (Å) | 3.34 | 3.60 |
| FSC threshold | 0.143 | 0.143 |
| Model resolution range (Å) | 1.9-40.4 | 2.2-10.2 |
| Map sharpening *B* factor (Å$^2$) | -50 | -150 |
| Model composition | | |
| Non-hydrogen atoms | 7245 | 7245 |
| Protein residues | 963 | 960 |
| Ligands | NAG:6, PPNDS:3 | NAG:6, PPADS:3 |
| *B* factors (Å$^2$) | | |
| Protein | 148.30 | 117.36 |
| Ligand | 189.65 | 146.40 |
| R.m.s. deviations | | |
| Bond lengths (Å) | 0.014 | 0.003 |
| Bond angles (°) | 1.257 | 0.561 |
| Validation | | |
| MolProbity score | 2.51 | 1.70 |
| Clashscore | 13.86 | 9.04 |
| Poor rotamers (%) | 3.70 | 1.22 |
| Ramachandran plot | | |
| Favored (%) | 93.50 | 97.17 |
| Allowed (%) | 6.39 | 2.83 |
| Disallowed (%) | 0.1 | 0 |
